# Age-trajectories of higher-order diffusion properties of major brain metabolites in cerebral and cerebellar grey matter using in vivo diffusion-weighted MR spectroscopy at 3T

**DOI:** 10.1101/2024.06.04.597406

**Authors:** Kadir Şimşek, Cécile Gallea, Guglielmo Genovese, Stephane Lehéricy, Francesca Branzoli, Marco Palombo

## Abstract

Healthy brain aging involves changes in both brain structure and function, including alterations in cellular composition and microstructure across brain regions. Unlike diffusion-weighted MRI (dMRI), diffusion-weighted MR spectroscopy (dMRS) can assess cell-type specific microstructural changes, providing indirect information on both cell composition and microstructure through the quantification and interpretation of metabolites’ diffusion properties. This work investigates age-related changes in the higher-order diffusion properties of three major intracellular metabolites (N-Acetyl-aspartate, Creatine and Choline) beyond the classical apparent diffusion coefficient in cerebral and cerebellar grey matter of healthy human brain. Twenty-five subjects were recruited and scanned using a diffusion-weighted semi-LASER sequence in two brain regions-of-interest (ROI) at 3T: posterior-cingulate (PCC) and cerebellar cortices. Metabolites’ diffusion was characterized by quantifying metrics from both Gaussian and non-Gaussian signal representations and biophysical models. All studied metabolites exhibited lower apparent diffusivities and higher apparent kurtosis values in the cerebellum compared to the PCC, likely stemming from the higher microstructural complexity of cellular composition in the cerebellum. Multivariate regression analysis (accounting for ROI tissue composition as a covariate) showed slight decrease (or no change) of all metabolites’ diffusivities and slight increase of all metabolites’ kurtosis with age, none of which statistically significant (p>0.05). The proposed age-trajectories provide benchmarks for identifying anomalies in the diffusion properties of major brain metabolites which could be related to pathological mechanisms altering both the brain microstructure and cellular composition.

## 1. Introduction

Healthy aging involves numerous and heterogeneous functional and structural changes in the brain depending also on the considered anatomical region. For instance, in-vivo studies showed that the cerebellum presents slower age-related morphological changes compared to the cerebral cortex (Liang & Carlson, 2020), possibly due to different microstructural properties. Indeed, the cerebellum contains 60 to 80% of the total amount of neurons in the brain for only 10% of the brain mass (Colin et al., 2001; Walløe et al., 2014). Investigating the neurobiological underpinnings of aging in the cerebellum is of interest as this structure projects to the entire brain and mediates cognitive functions affected by aging (Manto, 2022). Age-related changes have been shown in the cerebellum and cerebral cortices only at the macroscopic level by in-vivo studies, whereas microstructural changes have been mostly observed ex-vivo throughout life (Andersen et al., 2003), and in patients with diseases progressing with aging (Grimaldi & Manto, 2013; R. J. Louis et al., 2014). These studies showed different results, with loss of white matter (WM) up to 25% associated with loss of Purkinje and Granule cells (Andersen et al., 2003; Arleo et al., 2024) and thinning of dendritic trees of Purkinje cells (R. J. Louis et al., 2014).

Magnetic resonance imaging (MRI) studies have shown global macrostructural changes (volume loss) of grey matter (GM) and WM in the brain with aging (Andersen et al., 2003; MacDonald & Pike, 2021; Walhovd et al., 2005); cortical thinning in the cerebral cortex (Sowell et al., 2004) with prefrontal and frontal cortices (alongside hippocampus) most affected during aging (Jernigan et al., 2001); and loss of GM in the cerebellar cortex (Stalter et al., 2023).

Diffusion-weighted MR imaging (dMRI) is a powerful and widely used imaging tool to quantify human brain microstructure in-vivo and non-invasively (Alexander et al., 2019; Jones, 2010). Recent dMRI studies investigating variations of diffusion metrics with age observed a significant increase of mean diffusivity and decrease of fractional anisotropy in the cerebral cortex and subcortical regions (Helenius et al., 2002; Pfefferbaum et al., 2010; Raghavan et al., 2021; Schilling et al., 2022; Watanabe et al., 2013), while others remained inconclusive regarding the cerebellum (Behler et al., 2021; van Aalst et al., 2022).

Although very sensitive to microstructural changes, dMRI cannot unambiguously inform on changes in cellular composition due to the poor cell-type specificity of water molecules. In contrast, diffusion-weighted MRS (dMRS) provides higher cell-type specificity (Cao & Wu, 2017; Ligneul et al., 2023; Palombo et al., 2016, 2017; Palombo, Shemesh, et al., 2018; Ronen & Valette, 2015; Vincent et al., 2020), offering the opportunity to inform on alterations of both cellular composition and microstructure with age, through the interpretation of measurements of metabolite diffusion properties. Some of the major brain metabolites are purely intracellular (e.g., N-Acetyl-aspartate, NAA; creatine, Cr, and choline compounds, tCho) and cell-type specific (e.g., NAA mostly concentrated in neurons and tCho mostly concentrated in glia) and can be used to infer compartment specific microstructural changes (Ligneul et al., 2019, 2023; Palombo et al., 2016, 2017; Palombo, Ligneul, et al., 2018). A recent dMRS work showed that the apparent diffusion coefficient (ADC) of five major intracellular metabolites (myo-Inositol and Glutamate in addition to NAA, Cr and Cho) was faster in healthy older adults and depended on brain region, suggesting region-specific alterations in the intra-cellular microenvironment (Deelchand et al., 2020). However, it is still unknown how other informative diffusion properties of brain metabolites diffusion beyond the ADC change with aging. For example, the apparent diffusional kurtosis, a higher-order diffusion metrics that quantifies the degree of non-Gaussianity, could inform on the effect of restrictions and hinderance imposed by the microenvironment on the diffusion of intracellular metabolites (Jensen et al., 2005).

This work aims to fill this gap and provide first age-trajectories of higher-order diffusion properties of major intracellular metabolites (total N-acetyl-aspartate, tNAA: NAA + N-acetyl-aspartyl-glutamate, NAAG; tCho: glycero-phosphoryl-choline, GPC + phosphoryl-choline, PCho; and total creatine, tCr: Cr, + phospho-creatine, PCr) and to highlight potential microstructural changes with age in the cerebral and cerebellar GM using dMRS. We focused our investigation on cerebral and cerebellar cortices because of the role of cerebellum in mediating cognitive functions which are affected by aging in the brain and its higher microstructural complexity in contrast to cerebral cortex.

## 2. Material and Methods

### 2.1. Subjects

A cohort of twenty-five healthy adults consisting of 11 females and 14 males were recruited for this study. The age range of the participants spanned from 25 to 80 years, with a mean age of 50.2 years and a standard deviation of 20.2 years. Dividing the cohort into younger (<50 years) and older (>50 years) adults, we have 13 participants (6 females) with a mean age of 31.8 and a standard deviation of 7.1 years, and 12 participants (5 females) with a mean age of 70.2 and a standard deviation of 5.3 years, respectively. The healthy participant inclusion criteria involve absence of neurotropic treatment, psychiatric disorders, and cognitive function disorders. All subjects provided informed consent according to local procedures prior to the study. The study was approved by the local ethics committee.

### 2.2. Data acquisition and processing

dMRS data were acquired using a 3T Siemens Prisma scanner (Siemens Healthineers, Erlangen, Germany) with a 64-channel receive-only head coil at the Paris Brain Institute (Institut du Cerveau, ICM), France. Three-dimensional T_1_-weighted magnetization-prepared rapid gradient echo images (field of view, 256 (anterior – posterior) x 256 (foot – head) x 231 (right – left) mm^3^; isotropic resolution, 0.9 mm; repetition and echo time (TR/TE), 2300/2.08 ms; total acquisition time, 5 min. 17 sec. were acquired to position the spectroscopic region-of-interest (ROI) and to perform tissue segmentation. Two ROIs targeting GM in the cerebellum and posterior-cingulate-cortex (PCC) were examined using a diffusion-weighted semi-LASER sequence (Genovese, Marjańska, et al., 2021). The ROIs were defined as 5.3 cm^3^ (15×16×22 mm^3^) in the cerebellum and 8.0 cm^3^ (20×20×20 mm^3^) in the PCC to maximize GM volume fraction (above 70%) in both ROIs. Spectral data was recorded with a spectral bandwidth of 3000 Hz and complex data points of 2048 at TE of 125 ms. During measurements, pulse triggering was applied, setting the average TR to three cardiac cycles. Diffusion-weighting was applied using tetrahedral-encoding scheme in directions of (−1 −1 −1), (−1 1 1), (1 −1 1), and (1 1 −1). Six b-values (b = [0, 0.96, 3.85, 8.67, 15.41, 24.10] ms/µm^2^) were applied with an effective gradient duration (δ) of 26.4 ms (two pairs of bipolar gradients with 6.6 ms duration) and an effective diffusion gradient separation (Δ) of 62.5ms. The effective b-values were computed by including crusher and slice selection gradients as well as cross-terms compensation. Twenty-four transients were acquired for each diffusion-weighted condition and saved individually for further postprocessing. Water suppression was performed using variable power with optimized relaxation delays (VAPOR) and outer volume suppression (Tkac et al., 1999). The water suppression flip-angle was calibrated for each participant. Additionally, water signals were acquired using the same diffusion-weighted conditions for eddy-current correction, excluding ultra-high b-values due to poor water signal. B_0_ shimming was performed using a fast automatic shimming technique with echo-planar signal trains utilizing mapping along projections, FASTESTMAP (Gruetter & Tkac, 2000).

Spectral processing was performed by following the state-of-the-art guidelines (Ligneul et al., 2023) on MathWorks MATLAB R2022a (The MathWorks Inc., 2022). Zero-order phase fluctuations and frequency drifts were corrected on single transients before averaging using the NAA peak. A peak-thresholding procedure was applied, for each diffusion condition, to discard the transients with artefactual low signal-to-noise ratio (SNR) caused by non-translational tissue motion (Genovese, Marjańska, et al., 2021). After processing of the transients in an acquisition, e.g., a b-value measurement in a diffusion direction, the transients were averaged for independent data fitting.

GM, WM, and cerebrospinal fluid (CSF) volume fractions were calculated in the ROIs using the T1-weighted images and the segment tool of SPM12 and MATLAB routines.

### 2.3. Data fitting

For each diffusion-weighted condition, averaged spectra were fitted independently with LCModel (Provencher, 1993). The SNR of spectra was reported from LCModel’s output (i.e., the ratio between signal intensity at 2.01ppm and twice the root mean square of fit residuals).

The basis set was simulated with an in-house written routine in MATLAB based on the density matrix formalism (Henry et al., 2006) and using previously reported chemical shifts and *J*-couplings (Govindaraju et al., 2000; Kaiser et al., 2010). The basis set included ascorbate, aspartate, Cr, γ-aminobutyric acid, glucose, glutamate, glutamine, glutathione, GPC, *myo*-inositol, lactate, NAA, NAAG, PCr, PCho, phosphorylethanolamine, scyllo-inositol, and taurine. Independent spectra for the CH_3_ and CH_2_ groups of NAA, Cr, and PCr were simulated and included in the basis set.

### 2.4. Data analysis

To characterize the higher-order metabolites’ diffusion properties, multiple diffusion signal analyses were conducted including diffusion signal representations and biophysical models (Jensen et al., 2005; Ligneul et al., 2023; Palombo et al., 2017). The data and codes used to produce the results reported in this paper will be publicly available at https://github.com/kdrsimsek.

#### 2.4.1. dMRS signal representations

First, the direction-averaged diffusion signals were fitted monoexponentially up to *b*<5 ms/µm^2^ to estimate the apparent diffusion coefficient (*ADC*) and characterize Gaussian properties (Ligneul et al., 2023). Kurtosis signal representation (from Eq.5 in (Jensen et al., 2005)) was used to estimate the apparent diffusion kurtosis (*K*) and determine non-Gaussian properties of metabolites up to *b*<10 ms/µm^2^ (Genovese, Marjańska, et al., 2021).

#### 2.4.2. dMRS biophysical models

For biophysical modelling, the astro-sticks model was fitted to the direction-averaged signals at all b-values to estimate the apparent intra-stick axial diffusivity (*D*_*intra*_) (Ligneul et al., 2023; Panagiotaki et al., 2012)

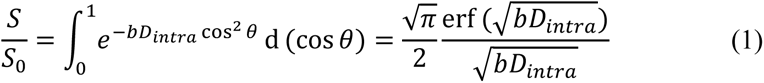

here, the equation describes direction-averaged diffusion signal for the astro-sticks model. *θ* is the angle between the main axis of a given stick and the applied diffusion gradient. **Error! Bookmark not** defined. Additionally, astro-sticks model was modified to incorporate an effective intra-stick axial diffusivity (*D*_*eff*_) defined as (Palombo et al., 2017; Palombo, Ligneul, et al., 2018; Sukstanskii & Yablonskiy, 2008; Yablonskiy & Sukstanskii, 2010):

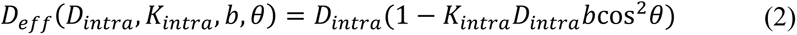

here, *K*_*intra*_is the apparent intra-neurite axial kurtosis and quantifies non-Gaussian diffusion characteristics stemming from hindering or restricting structures randomly displaced along the cellular processes, such as dendritic spines (Palombo, Ligneul, et al., 2018; Sukstanskii & Yablonskiy, 2008; Yablonskiy & Sukstanskii, 2010). The corresponding powder-averaged signal for the modified astro-sticks model is computed by numerical integration given in the following equation:

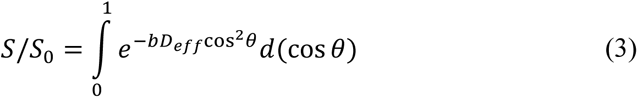

#### 2.4.3. Fitting routine

Data analysis was conducted within the Python programming environment. Following spectral quantification using LCModel, we estimated the diffusion-weighted signal amplitude from the area under each metabolites’ peak(s) and direction-averaged it at each b value to obtain the direction-averaged diffusion signal decay for each metabolite. Diffusion fitting was performed using Levenberg-Marquardt non-linear least squares optimization in the Python library ‘lmfit’ (https://pypi.org/project/lmfit/). No constraints were imposed on the modelling functions, but boundary conditions of each model parameter were defined to be positive and not to exceed free metabolites’ diffusivity 1.0 µm^2^/ms (Döring et al., 2018) and 3.0 for apparent kurtosis parameters (Jensen et al., 2005). Three major metabolites were examined: tNAA as a neuronal biomarker; tCho as a glial biomarker; and tCr as a biomarker comprised in both neuronal and glial cells. Notably, one direction-averaged diffusion signal in the cerebellum, acquired from a subject, suffered from very poor SNR; hence, excluded from the diffusion analysis.

### 2.5. Statistical Analysis

Linear regression was performed on all estimated parameters to determine age-trajectories with computed 95% confidence interval and prediction limits. To analyze the specific impact of age on the changes of diffusion metrics, a regression analysis with age as the independent variable and each estimated model parameter as the dependent variable was performed, also accounting for 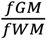 (the ratio between GM and WM volume fractions) as covariate by fitting the following expression: *y* ∼ *β* _0_ + *β* _1_ ⋅ *age* + *β* _2_ ⋅ *fGM*/*fWM*.

Additionally, an independent T-test between younger (age < 50) and older (age≥ 50) people was performed to assess statistically significant differences between younger and older adult groups. Bonferroni correction was applied for only T-test, including two brain regions and three metabolites for each diffusion metric and the p-value, the threshold for statistical significance, was redefined to be 0.0083 (0.05/6). In both statistical analyses, the parameters values converging to the lower bound in the fitting were excluded from the age-trajectory analysis because considered unreliable.

## 3. Results

To simplify inspection of the findings, a color-coding scheme is used to identify the cerebellum and the PCC results as blue and red more clearly, respectively.

Exemplary diffusion-weighted spectra acquired from both brain regions are shown in **Figure 1A** which exhibit good spectral quality – linewidths at b_0_/b_max_: 4.17/4.84 Hz in the cerebellum and 3.30/4.67 Hz in the PCC. SNRs obtained from the corresponding LCModel fit results were 18±3 and 24±4 (mean ± standard deviation over all subjects) at *b* = 0 (i.e., no diffusion-weighting) and 7±2 and 6±2 at the highest *b* value in the cerebellar and cerebral cortexes, respectively. The tissue volume fractions (mean ± standard deviation over all subjects) were as follows: fGM: 0.82 ± 0.05 (GM volume fraction), fWM: 0.12 ± 0.05 (WM volume fraction), and fCSF: 0.06 ± 0.03 (CSF volume fraction) in the cerebellum; fGM: 0.69 ± 0.07, fWM: 0.14 ± 0.03, and fCSF: 0.17 ± 0.08 in the PCC. The localizations of spectroscopic voxels in both ROIs are depicted in **Figure 1B**. Furthermore, the age-trajectory for 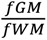 ratio was investigated for variations with age and reported in **Figure 1C**. A significant decrease with age in 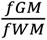 in the cerebellum was observed, while no significant change in the PCC.

**Figure 1:**
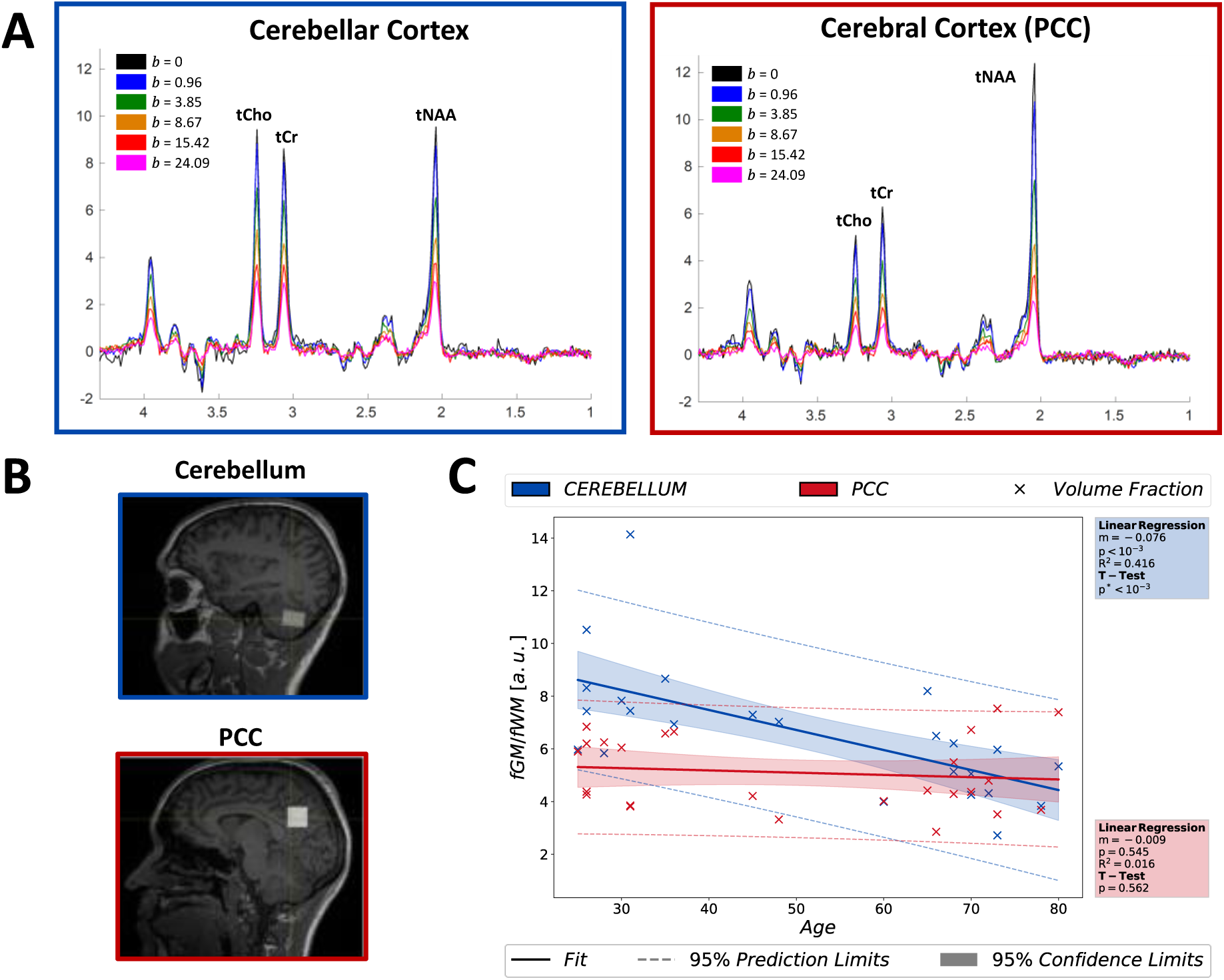
(**A**) Diffusion-weighted spectra are illustrated for both regions of interest; cerebellum (blue frame, left) and PCC (red frame, right). Direction averaged dMRS signals exhibit excellent spectral quality. Color-coding in the legends displays b-values in the units of ms/μm^2^. (**B**) Regions of interest are demonstrated on T_1_-w images. (**C**) Age-trajectories of fGM/fWM ratio in both ROIs and the results of statistical analyses reporting only a significant decrease in fGM/fWM in the cerebellum with age. (p*<0.00833 indicates statistical significance for all tests). Abbreviations: PCC, posterior cingulate cortex; fGM, grey matter volume fraction; fWM, white matter volume fraction; tNAA, total N-Acetyl-aspartate; tCho, total choline; tCr, total creatine; ROI, region of interest.

The Cramer-Rao lower bound (CRLB) obtained from LCModel fit was used to assess the quality of the quantification. Overall fit results are excellent with low CRLBs (<5%) in both ROIs for the non-diffusion weighted spectra. Estimated CRLBs for the highest *b* values were averaged over different diffusion directions and yielded CRLB_tNAA_=4%, CRLB_tCho_=6%, and CRLB_tCr_=4% in the cerebellum and CRLB_tNAA_=7%, CRLB_tCho_=10%, and CRLB_tCr_=5% in the PCC. In addition to CRLB, no other exclusion criteria were employed (Kreis, 2016).

### 3.1. Metabolite Diffusion Properties

Metabolite diffusion signals obtained from all subjects are displayed in **Figure 2A** for both cerebellum and PCC. The diffusion signals obtained from all participants (light) are reported alongside the corresponding cohort averages (dark). Overall, slower metabolite diffusion was observed for all metabolites in the cerebellum compared to PCC. **Figure 2B** presents the results of the estimated diffusion parameters from all subjects as a box-whiskers plot for all signal representations and biophysical models. The corresponding mean values of the estimated parameters obtained from the cohort are charted in **Table 1**. In the cerebellum, the model parameters for one dataset could not be estimated (and highlighted as an outlier with values of zero), due to low SNR at higher b-values. In all cases, the estimated apparent diffusivities (*ADC*s & *D*_*intra*_) are lower in the cerebellum than in the PCC. Correspondingly, the kurtosis estimates (*K* and *K*_*intra*_) are higher in the cerebellum than in the PCC, for all metabolites. Noticeably, *K*_*intra*_ of tCho and tCr in both ROIs exhibit high variability due to relatively higher CRLB; e.g. in the tCho results, the median values in each metabolite result is at the lower bound while the mean values are higher as shown in **Figure 2B**.

**Figure 2:**
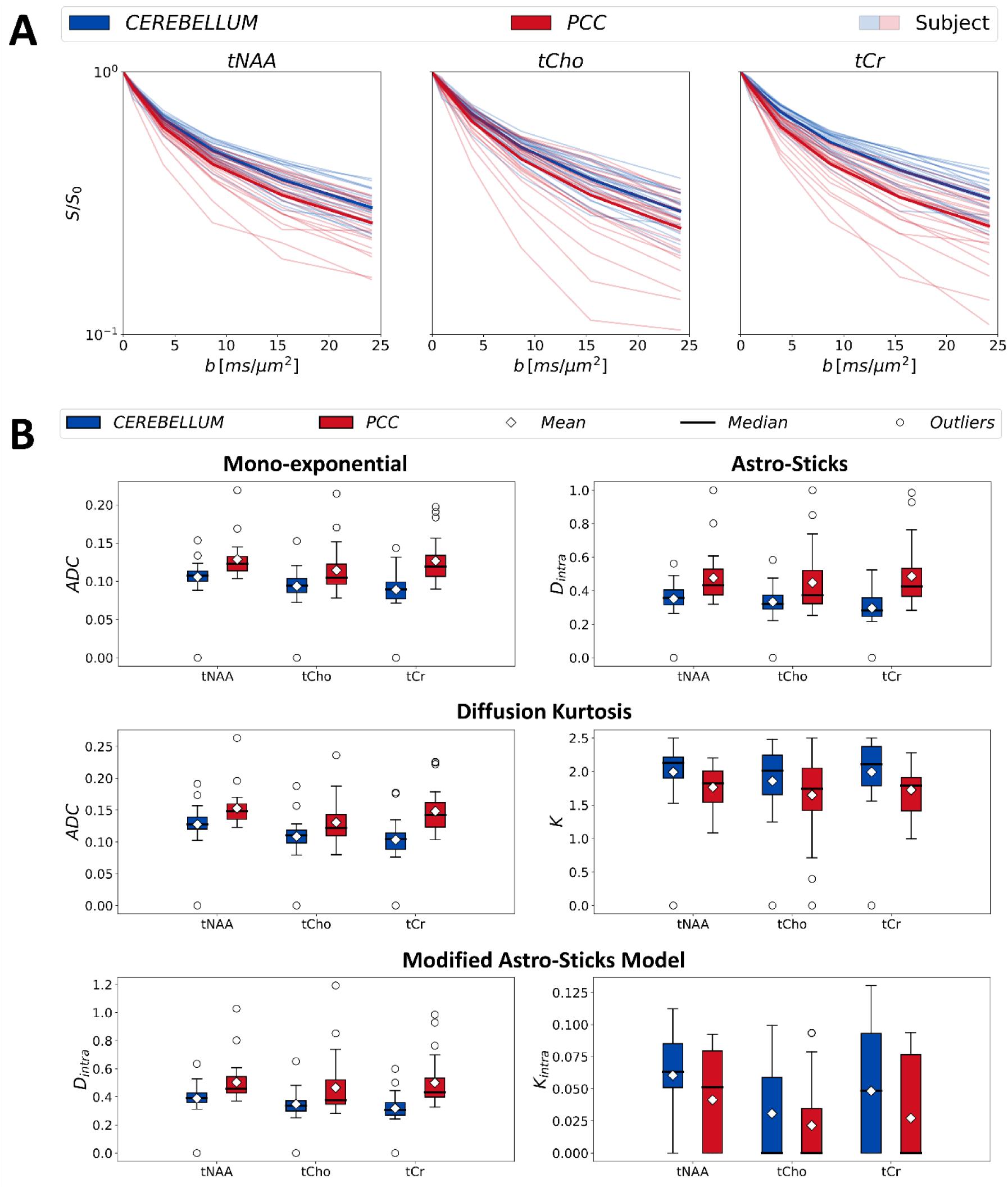
(**A**) Diffusion signals of tNAA, tCho, and tCr obtained from each subject (light) and cohort averaged signals (dark) are illustrated in the figure for both brain regions: cerebellum (blue) and posterior-cingulate-cortex (red). (**B**) The estimated parameters of each metabolite by mono-exponential, kurtosis representations and astro-sticks and modified astro-sticks models from each subject are illustrated in the box-and-whiskers plot for both region of interests: cerebellum (blue) and PCC (red). Abbreviations: PCC, posterior cingulate cortex; tNAA, total N-Acetyl-aspartate; tCho, total choline; tCr, total creatine; ADC: apparent diffusion coefficient; K, apparent diffusion kurtosis; *D*_*intra*_, apparent intra-neurite axial diffusivity; *K*_*intra*_, apparent intra-neurite axial.

**Table 1:** Estimated model parameters obtained from cohort averaged diffusion signals are charted with the corresponding error values in the fit (estimation ± error). All signal representations and biophysical model results are tabulated in the table. “ ̶ “ indicates that the estimations converge to zero. Abbreviations: PCC, posterior cingulate cortex; tNAA, total N-Acetyl-aspartate; tCho, total choline; tCr, total creatine; ADC: apparent diffusion coefficient; K, apparent diffusion kurtosis; *D*_*intra*_, apparent intra-neurite axial diffusivity; *K*_*intra*_, apparent intra-neurite axial kurtosis.

| ROI | Fit | Parameter | tNAA | tCho | tCr |
| --- | --- | --- | --- | --- | --- |
| CEREBELLUM | Monoexp | $ADC$ | $0.109 \pm 0.007$ | $0.097 \pm 0.004$ | $0.092 \pm 0.004$ |
| | Kurtosis | $ADC$ | $0.132 \pm 0.006$ | $0.112 \pm 0.003$ | $0.107 \pm 0.003$ |
| | | $K$ | $2.112 \pm 0.108$ | $2.000 \pm 0.083$ | $2.204 \pm 0.078$ |
| | Astro-sticks | $D_{intra}$ | $0.360 \pm 0.012$ | $0.340 \pm 0.007$ | $0.300 \pm 0.007$ |
| | Modified Astro-sticks | $D_{intra}$ | $0.400 \pm 0.016$ | $0.340 \pm 0.007$ | $0.326 \pm 0.010$ |
| | | $K_{intra}$ | $0.071 \pm 0.014$ | — | $0.064 \pm 0.018$ |
| PCC | Monoexp | $ADC$ | $0.127 \pm 0.006$ | $0.112 \pm 0.002$ | $0.126 \pm 0.003$ |
| | Kurtosis | $ADC$ | $0.152 \pm 0.002$ | $0.130 \pm 0.001$ | $0.148 \pm 0.001$ |
| | | $K$ | $1.778 \pm 0.034$ | $1.733 \pm 0.026$ | $1.747 \pm 0.020$ |
| | Astro-sticks | $D_{intra}$ | $0.460 \pm 0.007$ | $0.422 \pm 0.018$ | $0.463 \pm 0.006$ |
| | Modified Astro-sticks | $D_{intra}$ | $0.488 \pm 0.009$ | $0.422 \pm 0.045$ | $0.463 \pm 0.023$ |
| | | $K_{intra}$ | $0.047 \pm 0.010$ | — | — |

### 3.2. Age-trajectories

The age-trajectories for apparent diffusivities (*ADC* & *D*_*intra*_) of monoexponential representation and astro-sticks model are grouped together and presented in **Figure 3**. The apparent diffusivities presented similar trends with age for all metabolites: increasing in the PCC and decreasing in the cerebellum.

**Figure 3:**
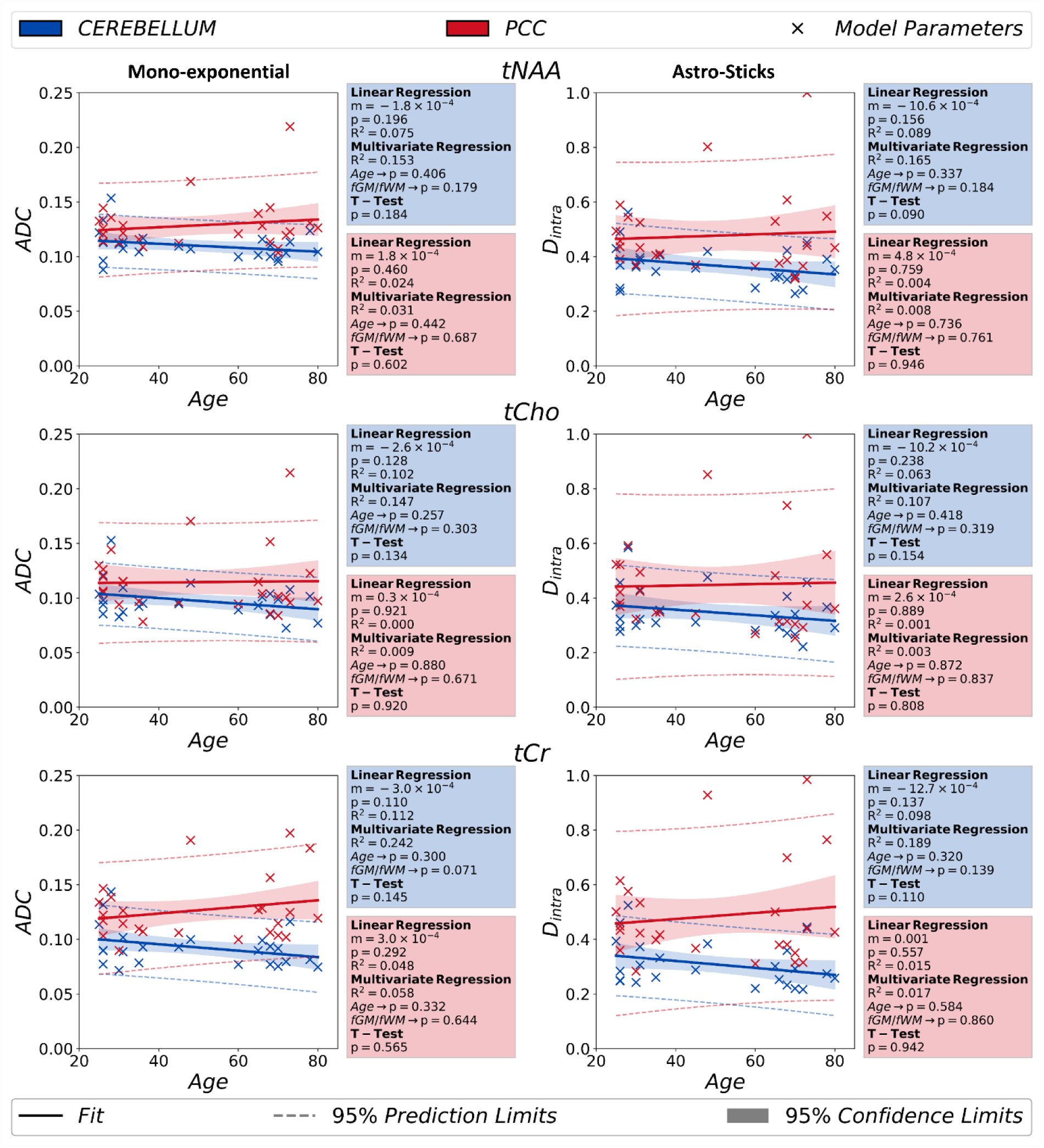
The results obtained from monoexponential signal analysis (*b*<5ms/μm^2^) (*ADC*) and astro-stick model (*D*_*intra*_) are documented in the figure. The independent T-test analyses performed between younger and older groups do not report any statistically significant change in these parameters with aging. The p-value in linear regression is a measure for how significant the estimated slope is in the analysis. (p*<0.00833 indicates statistical significance for the T-Test) Abbreviations: PCC, posterior cingulate cortex; fGM, grey matter volume fraction; fWM, white matter volume fraction; tNAA, total N-Acetyl-aspartate; tCho, total choline; tCr, total creatine; ADC: apparent diffusion coefficient; *D*_*intra*_, apparent intra-neurite axial diffusivity

The age-trajectories for diffusion kurtosis (*ADC* & *K*) and modified astro-sticks model (*D*_*intra*_& *K*_*intra*_) analyses were grouped together and showed in **Figure 4**. The age-trajectories for the diffusion kurtosis parameters depicted in **Figure 4A** predominantly show similar trends for all metabolites except for the *ADC* of tCr in the PCC and *K* of tCho in the cerebellum, which exhibit opposite trend. Likewise, the age-trajectories of modified astro-sticks model parameters show decreasing trend in *D*_*intra*_ and increasing trend in *K*_*intra*_for all metabolites in both ROIs as illustrated in **Figure 4B**. Only exception is the *D*_*intra*_of tCr in the PCC showing increasing trend.

**Figure 4:**
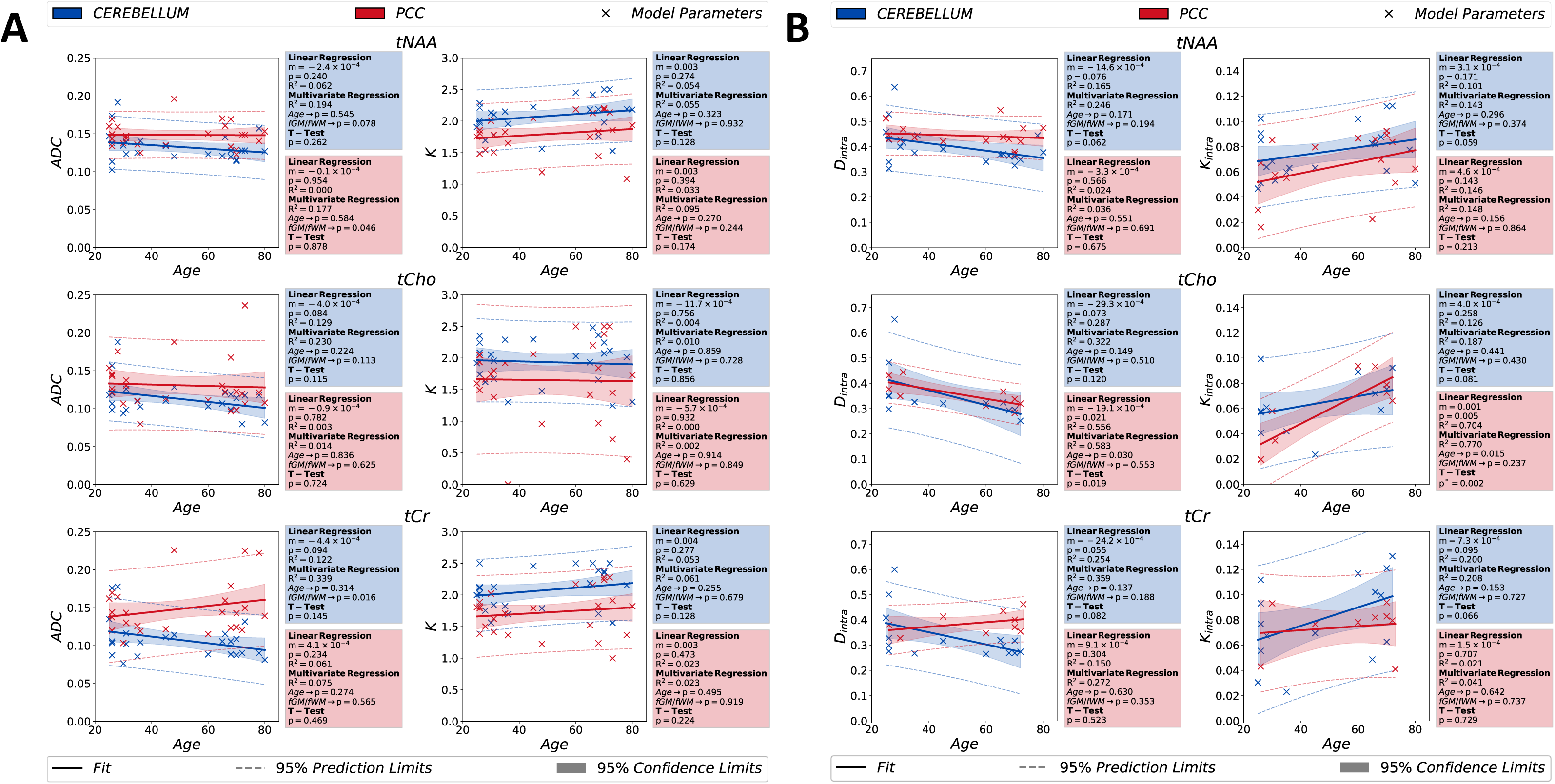
Age dependences of the estimated model parameters for kurtosis (***ADC*** & ***K***) in (A) and modified astro-stick model (***D*_*intra*_**& ***K*_*intra*_**) in (B), obtained from studied metabolite signals, are depicted in the figure. For each brain region, a linear regression, a regression analysis using age and fGM/fWM as independent and dependent variables, respectively, and a paired T-test between two groups [age < 50 and age ≥ 50] are performed to analyze impact of age and tissue composition on the estimated parameters. For statistical tests, the confidence and prediction limits are also depicted in the figure. (p*<0.00833 indicates statistical significance for the T-test) Abbreviations: PCC, posterior cingulate cortex; fGM, grey matter volume fraction; fWM, white matter volume fraction; tNAA, total N-Acetyl-aspartate; tCho, total choline; tCr, total creatine; ADC: apparent diffusion coefficient; K, apparent diffusion kurtosis; ***D*_*intra*_**, apparent intra-neurite axial diffusivity; ***K*_*intra*_**, apparent intra-neurite axial kurtosis

Overall, the statistical analyses performed over diffusion metrics of tNAA (the neuronal biomarker), tCho (glial biomarker) and tCr (less cell-type specific) do not report any significant change with age for all the higher-order diffusion metrics investigated in this study (p>0.05). Notably, the T-test results of tCho *K*_*intra*_show only significant increase in the PCC. Considering the high noise level in tCho and the median value of tCho *K*_*intra*_at the lower bound in the modified astro-sticks model fitting, this outcome needs to be treated carefully.

## 4. Discussion

This work investigates variations in the higher-order diffusion properties of major intracellular brain metabolites with healthy aging in the cerebral and cerebellar GM in-vivo in human brain using dMRS and clinical 3T MRI scanner.

### 4.1. Metabolites apparent diffusivity in cerebellar and cerebral GM

Apparent diffusivities (*ADC* & *D*_*intra*_) of the studied metabolites agree with literature findings (Branzoli et al., 2014; Deelchand et al., 2018; Döring et al., 2018; Döring & Kreis, 2019; Ingo et al., 2018; Kan et al., 2012; Najac et al., 2016; Palombo et al., 2016; Şimşek et al., 2022). Relatively slower metabolite apparent diffusivities in the cerebellum might stem from higher microstructural complexity of cellular composition compared to PCC: the Purkinje and granule cells are highly abundant in the cerebellum (E. D. Louis et al., 2014) while the PCC is comprised by the less complex Pyramidal neurons.

### 4.2. Age-dependence of metabolites apparent diffusivity

Overall, estimated apparent diffusivities did not present any significant trend nor changes with age, in contrast to mono-exponential ADCs reported by the only study in the literature (Deelchand et al., 2020). This difference might originate from having different sample size and more likely from different ROI tissue volume composition. In contrast to our work, Deelchand et al. recruited more participants and in two age groups (young: 18-22 and old: 70-83 years old). Regarding distinctions in tissue composition, the WM content in the PCC ROI in our work is around half that in the Deelchand’s work (our work, fWM=14%; Deelchand’s work, fWM≃30%). Higher fGM in our ROIs leads to a more isotropic microenvironment for metabolite diffusion; thus, a weaker dependence of metabolite apparent diffusivity on the fiber orientation. Other contributing factors might be differences in diffusion times (62.5 in our work and 125 ms in Deelchand’s work) and encoding schemes. Previous studies have shown that diffusion times have strong effects on estimated ADCs in both GM and WM (Assaf & Cohen, 1998; Döring & Kreis, 2019; Ligneul et al., 2017; Ligneul & Valette, 2017), while TE-dependence of metabolites ADC was only significant in ROIs with high content of WM (Branzoli et al., 2014) like in Deelchand’s work and was negligible for ROIs with high content of GM (Ligneul et al., 2017), like in our study.

### 4.3. Metabolites apparent kurtosis and non-Gaussianity in cerebellar and cerebral GM

Estimated metabolite diffusion kurtosis *K* values agree with current literature (Döring et al., 2023; Genovese, Marjańska, et al., 2021; Ingo et al., 2018; Mougel et al., 2023). In accordance with metabolite apparent diffusivities, the *K* & *K*_*intra*_ for all metabolites in the cerebellum compared to PCC agree with the expected higher complexity of the cellular microenvironment. For instance, the Purkinje cells in the cerebellum have higher spine density and higher branching order (Santamaria et al., 2006) in contrast to the PCC, which comprises mostly Pyramidal cells with lower spine density and branching order (Holtmaat et al., 2005). Therefore, the higher microstructural complexity in the cerebellum might lead to higher tNAA (the neuronal biomarker) apparent *K* & *K*_*intra*_. Additionally, the relatively higher *K* & *K*_*intra*_ values for tCho (glial biomarker) in the cerebellum might be due to the presence of highly arborized Bergmann glia (Sild & Ruthazer, 2011). The same rationale can explain the observed lower diffusivities in the cerebellum compared to the PCC.

### 4.4. Age-dependence of metabolites apparent kurtosis and non-Gaussianity

The age-trajectories of metabolite diffusion properties reveal overall similar trends for apparent diffusivities (*ADC* & *D*_*intra*_) from signal representations and biophysical models. The significant increase with age in *K*_*intra*_ of tCho in the PCC requires cautious interpretation due to the low SNR and high CRLB in the cerebellum data. **Figure 2B** illustrates that the median value of tCho *K*_*intra*_ is at the lower bound. Therefore, the low SNR in tCho might cause instability in fitting of modified astro-sticks model that resulted in a significant increase in *K*_*intra*_ in the PCC.

### 4.5. Analysis of potential confounders: the negligible impact of ROI tissue composition

The multivariate regression analysis does not report any significant impact of the accounted variables age (as independent) and *fGM*/*fWM* (as dependent) on the variation of diffusion metrics (p>0.05). Hence, the trend in age-trajectory cannot be attributed to changes in the volume fractions of tissue compositions in the ROIs. The only exception is for the tCho *K*_*intra*_in the PCC (**Figure 4B**), having p-value for age just below the threshold (p=0.013). However, the observed change possibly arises from the encountered model fitting issues in tCho, the glial biomarker.

A previous study reported that the ADC of tNAA changes by 8% between young and old groups and argued that the contribution stemming from their ROIs tissue composition would be relatively small in comparison to the observed percentage change in the tNAA ADC (Deelchand et al., 2020). A similar argument can be made in our study. For instance, *D*_*intra*_ of tCr from the astro-sticks model exhibits the strongest change of about 10% increase within the age limit in the PCC (**Figure 4B**). However, the change in the ROI tissue composition, 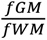, is only around 2% (**Figure 1C**) and cannot alone explain the changes observed in the PCC. Moreover, the multivariate analysis of the corresponding age-trajectory does not demonstrate any dependence on 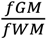 (p>0.05). Therefore, other factors, more directly linked to changes in the tissue microstructure and cellular composition might explain the observed trends in age-trajectories. A longitudinal study monitoring microstructural alterations in stroke linked an increase of tCr ADC with astrogliosis and glial reactivity in the presence of neuroinflammation in stroke patients (Genovese et al., 2023). Accordingly, an increase in astrogliosis and glial reactivity with aging was also reported in the literature (Cotrina & Nedergaard, 2002) that might explain the slight increasing trend in the *D*_*intra*_ of tCr.

### 4.6. Limitations

Our study has a few limitations that future studies may want to address. Diffusion-weighted spectra are very sensitive to the bulk or physiological motion occurring during the acquisition, causing variations in signal amplitude and phase (Branzoli et al., 2014; Döring et al., 2018; Ligneul et al., 2023; Şimşek et al., 2022). Employing cardiac triggering during measurements and performing SNR thresholding partially eliminated these (Genovese, Marjańska, et al., 2021; Ligneul et al., 2023). Due to poor water signal at high b-values, eddy-current correction was not applied to the spectra acquired at ultra-high b values. Because of the relatively small sample size, the statistical analysis was performed on two age groups (age<50 and age≥50) to accommodate enough datasets.

### 4.7. Importance and potential impact

The age-trajectories here reported are a precious resource for the community because they provide reference values for a large set of diffusion properties in two brain regions of potential interest for many diseases (e.g., Alzheimer’s disease and motor disorders), previously unavailable. As an example, choline compound is known as a neuroinflammation biomarker (De Marco et al., 2022; Genovese, Palombo, et al., 2021; Lind et al., 2021). A recent dMRS study (De Marco et al., 2022; Genovese, Palombo, et al., 2021) showed a significant increase in tCho ADC in the thalamus with neuroinflammation. The age-trajectories reported here provide reference values for the healthy brain cerebellum and PCC, suggesting that the age-related changes of tCho ADC are less than 10% (decrease in the cerebellum and increase in the PCC with age) which can help further interpreting tCho diffusivity results in studies of neuroinflammation in these brain regions. Age is often found to be a significant covariant in the analyses of the changes of metabolites diffusivity. Here we show to what extent age indeed alter the diffusion properties of major metabolites in ROIs mostly comprised of GM (> 70%). For instance, for the widely used ADC index, no statistically significant changes are observed for tNAA, tCr and tCho between younger (<50) and older (≥50) adults, with metabolites ADCs being overall less than 10% lower in older adults in the cerebellum, and less than 5% higher in older adults in the PCC.

## 5. Conclusion

This study offers previously unavailable age-trajectories of major intracellular brain metabolites’ diffusion properties in cerebral and cerebellar GM. We showed that observed variations in metabolite diffusion properties with healthy aging are minimal and most likely caused by age-related microstructural changes, demonstrating the potential utility of the metabolites high-order diffusion parameters as new (neuronal and glial) biomarkers of tissue pathology. The proposed age-trajectories provide benchmarks for identifying anomalies in the diffusion properties of major brain metabolites, which could be related to pathological mechanisms altering both the GM microstructure and cellular composition.

## Research Funding State

- This work, Kadir Şimşek and Marco Palombo are supported by UKRI Future Leaders Fellowship (MR/T020296/2).
- This project has received funding from Engineering and Physical Sciences Research Council (EPSRC EP/N018702/1). FB, CG and SL acknowledge support from the programs ‘Institut des neurosciences translationnelle’ ANR-10-IAIHU-06 and ‘Infrastructure d’avenir en Biologie Santé’ ANR-11-INBS-0006.

## Acknowledgements

• The authors would like to thank Dr. Edward J. Auerbach and Dr. Małgorzata Marjańska for providing us with the dMRS sequence for the Siemens platform and simulating the basis set for spectral fitting.

## CRediT authorship contribution statement

Kadir Şimşek: Conceptualization, Formal analysis, Methodology, Investigation, Writing - original draft; Cécile Gallea: Writing-review & editing; Guglielmo Genovese: Methodology, Formal analysis; Data curation, Writing-review & editing; Stephane Lehéricy; Writing-review & editing; Francesca Branzoli: Formal analysis; Funding acquisition, Methodology, Project administration, Writing - review & editing; Marco Palombo: Conceptualization, Methodology, Supervision, Resources, Project administration, Investigation, Writing - review & editing

